# Heritable changes in chromatin contacts linked to transgenerational obesity

**DOI:** 10.1101/2022.10.27.514145

**Authors:** Richard C. Chang, Riann J. Egusquiza, Yikai Huang, Angélica Amorim Amato, Erika M. Joloya, Hailey B. Wheeler, Angela Nguyen, Keiko Shioda, Junko Odajima, Toshi Shioda, Bruce Blumberg

**Affiliations:** Department of Developmental and Cell Biology, University of California, Irvine, CA, USA, 92697-2300; Department of Pharmaceutical Sciences, University of California, Irvine, CA, USA, 92697-2300; Department of Biomedical Engineering, University of California, Irvine, CA, USA, 92697-2300; Department of Pharmaceutical Sciences, University of Brasilia, Brasilia, Brazil; Center for Cancer Research, Massachusetts General Hospital, Bldg 149, 13th Street, Charlestown, MA, 02129, USA

## Abstract

Burgeoning evidence demonstrates that responses to environmental exposures can be transmitted to subsequent generations through the germline without DNA mutations^1,2^. This is controversial because underlying mechanisms remain to be identified. Therefore, understanding how effects of environmental exposures are transmitted to unexposed generations without DNA mutations is a fundamental unanswered question in biology. Here, we used an established murine model of transgenerational obesity to show that direct or ancestral exposure to the obesogen tributyltin (TBT) elicited persistent changes in topologically associating domains (TADs) in primordial germ cells (PGCs) isolated from embryos of exposed and subsequent unexposed generations. New TAD boundaries were formed within the *Ide* gene encoding insulin degrading enzyme in the exposed PGCs, then stably maintained in PGCs of the subsequent (unexposed) two generations. Concomitantly, *Ide* mRNA expression was decreased in livers of male descendants from the exposed dams. These animals were hyperinsulinemic and hyperglycemic, phenocopying *Ide*-deficient mice that are predisposed to adult-onset obesity. Creation of new TAD boundaries in PGCs, suppression of hepatic *Ide* mRNA, increased fat mass, hyperinsulinemia and hyperglycemia were male-specific. Our results provide a plausible molecular mechanism underlying transmission of the transgenerational predisposition to obesity caused by gestational exposure to an environmental obesogen. They also provide an entry point for future studies aimed at understanding how environmental exposures alter chromatin structure to influence physiology across multiple generations in mammals.

---

Several possible mechanisms have been proposed to explain transgenerational inheritance^2^. These included epigenetic alterations such as DNA methylation^1^, histone methylation^3^, histone retention^4,5^, and the transmission of small, non-coding RNAs^6^. Each has some merit for transmitting the results of direct exposure. However, none provides a mechanistic understanding for how phenotypes can be transmitted to unexposed descendants^2^. DNA methylation is erased genome-wide twice each generation in mammals^7^. This makes it unclear how the large number of differentially methylated regions we previously observed in white adipose tissue (WAT) of male F4 tributyltin (TBT)-group mice^8^ could have been transmitted to an unexposed generation. How histone methylation at specific loci can be transmitted across generations and why only some histones are retained during spermatogenesis are unknown. It remains uncertain how expression of non-coding RNAs, such as those that were reported in F1 sperm and seminal fluid^9^ could be transmitted to multiple future generations. Our preceding studies suggested that stable alterations of higher-order chromatin structures might provide a unifying theory to explain mammalian transgenerational inheritance^8,10^. Support for such a model requires the identification of persistently altered regions of higher-order chromatin structure. Therefore, we sought to identify regions of persistently altered chromatin structure in PGCs from mice exposed to TBT (F1) and unexposed (F2, F3) generations and associate these specific changes with the transgenerational obesity phenotypes observed.

## Ancestral TBT exposure led to a transgenerational predisposition to increased WAT mass

In the new transgenerational experiment 4 (T4) detailed here, effects of ancestral TBT exposure throughout gestation were confirmed to be much stronger in male versus female F2 and F3 generation C56BL/6J mice. This confirms our previous transgenerational experiments denoted as T1^11^, T2^8^ and T3^12^ and another study using OG2 C56BL/6J mice^13^. Salient findings included increased WAT depot size, more overall body fat and in some cases, increased body weight (Fig. 1). Male F2 animals ancestrally exposed to TBT accumulated significantly more WAT after the diet was changed from a standard chow diet (SD) to a higher fat diet (HFD) at 5 weeks of age (Fig.1a); body weight did not differ between groups. In contrast, F3 male animals did not show increased body weight or fat mass when switched to the HFD at 5 weeks (Extended data Fig.1). A small increase in body fat began at 11 weeks, but never reached statistical significance (Extended data Fig.1). We hypothesized that HFD challenge had started too soon compared with previous experiments. Therefore, we switched sibling DMSO- and TBT-group F3 animals that had been maintained on the SD to HFD at 17 weeks of age. TBT group males rapidly accumulated body weight and fat mass compared with controls; these differences became statistically significant at 19 (body fat) or 20 weeks (body weight) (Fig. 1b). No effects of ancestral TBT exposure on fat accumulation were observed in females (Fig. 1c, d).

**Fig. 1.**
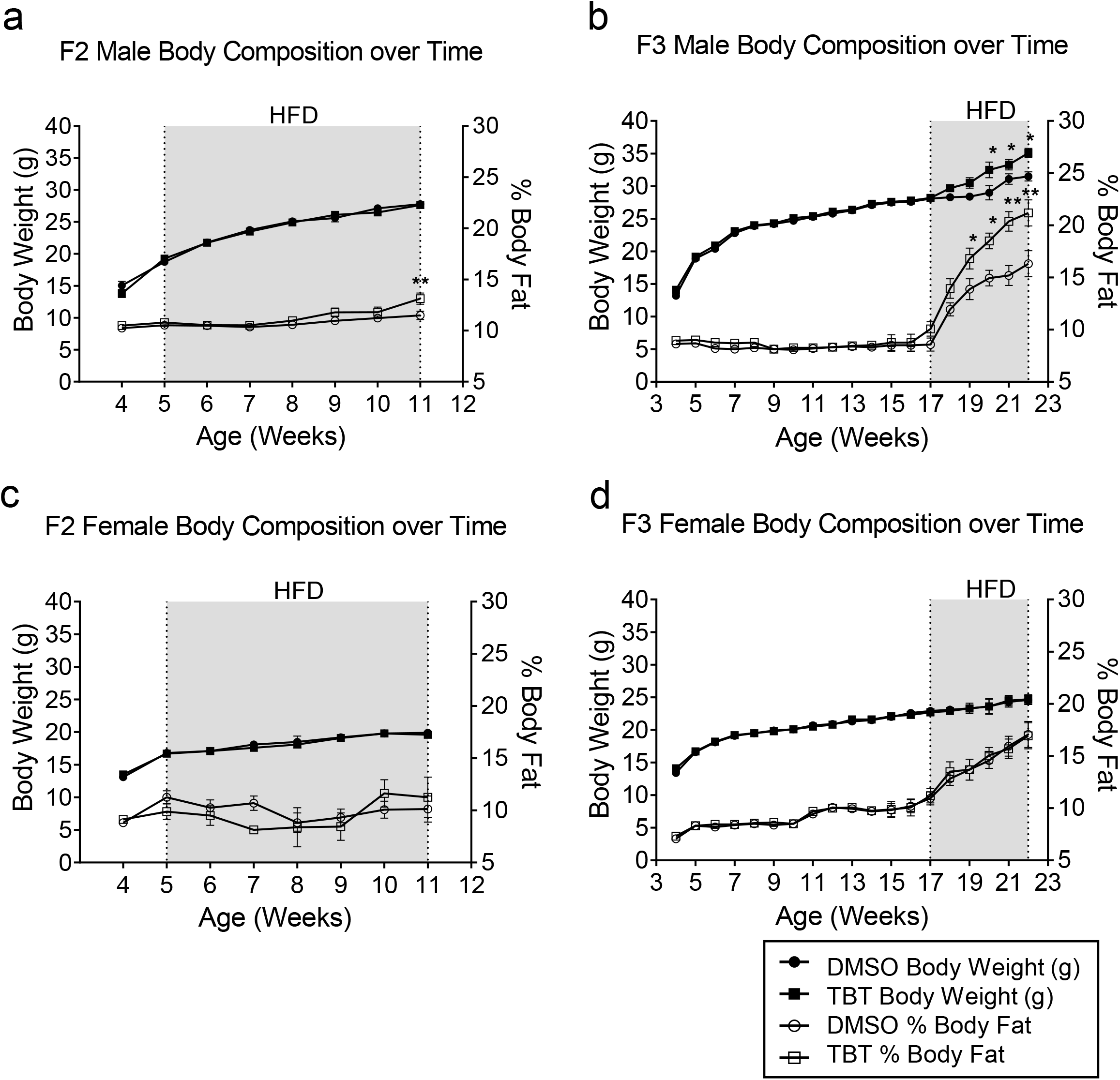
Mice ancestrally exposed to TBT exhibited increased fat content in F2 and F3 male descendants. Body weight and relative body composition of **a**, F2 male descendants (n = 15), **b**, F3 male descendants (n = 16), **c**, F2 female descendants (n = 17), and **d**, F3 female descendants (n = 16) throughout the time course of the experiment. Gray area indicates the period of diet challenge. Statistical significance was determined using two-way ANOVA. Pair-wise Bonferroni post-hoc tests were used to compare different groups. Data are presented as mean ± s.e.m. *p < 0.05; **p < 0.01.

## Ancestral TBT exposure caused transgenerationally stable chromatin contact alterations in the genome of male PGCs

To determine whether chromatin structure was stably altered after ancestral TBT exposure, we performed Hi-C seq of PGCs isolated from E13.5 embryonic gonads of F1-F3 mice, pooled by litter and sex. Approximately 10,000 – 15,000 FACS-enriched PGCs were subjected to Hi-C sequencing, data generation and analysis (Extended data Fig. 2). Successful detection of a known set of TADs around the *HoxD* gene cluster^14^ demonstrated the validity of our Hi-C seq data (Extended data Fig. 2h, i)

PGCs in F1 embryos were directly exposed to TBT while the embryos were within the treated F0 dams. Those isolated from F2 or F3 embryos were not exposed. PGCs in F2 embryos became gametes producing F3 animals, which showed a transgenerational predisposition to diet-induced obesity^8,11-13^. The global profiles of chromatin contacts were well conserved in PGC genomes across sex, F0 exposure to TBT, or F1-F3 generations (Extended data Fig. 3a). The median distance between two chromatin contacts was approximately 1 megabase (Extended data Fig. 3b), which agrees with the previously reported size of TADs in the mouse genome (880 kb)^15^.

Genome-wide assessments of altered chromatin contacts in male PGCs in F1 embryos revealed significant gains in TAD boundaries associated with binding sites for the canonical loop stabilizing complex (LSC) (Fig. 2a) components CTCF, cohesin (including SMC3), and RAD21 (Fig. 2b). Male PGCs in F2 embryos gained TAD boundaries associated with binding sites for RXRA, TRIM28, and pluripotency transcription factors POU5F1 and NANOG in addition to LSC (Fig. 2b). Male PGCs in F3 embryos lost the gained TADs associated with LSC. Although female PGCs in F1 and F2 embryos also gained some TADs associated with CTCF, formation of LSC might have been incomplete as concomitant association of SMC3 or RAD21 was not detected (Fig. 2b).

**Fig. 2.**
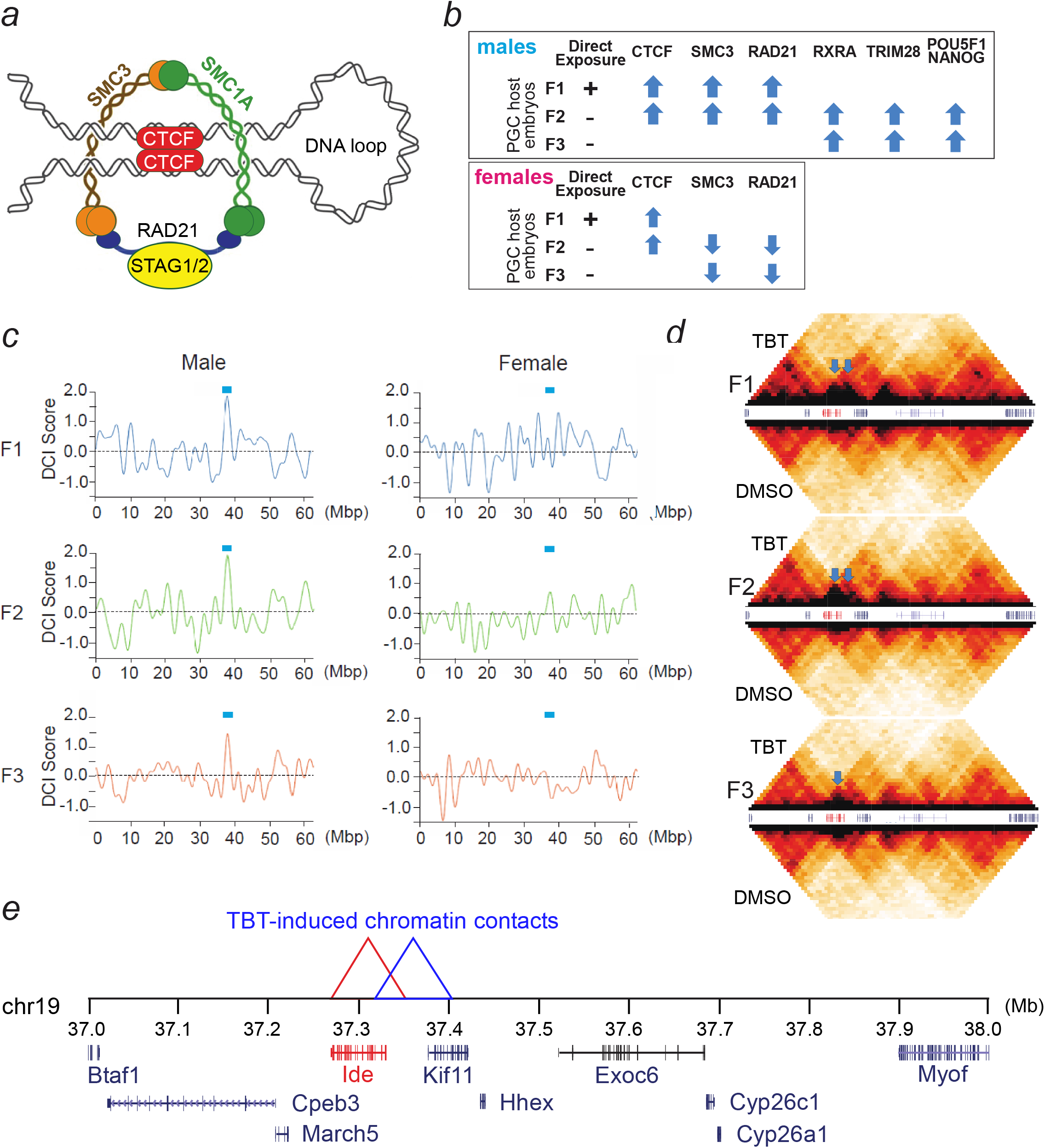
Transgenerationally transmitted differential chromatin contacts (DCC). **a**, Schematic representation of the CTCF/cohesion complex at the boundary of a DNA loop. **b**, results of BART-3D analysis of association between differential TADS and known transcription factor binding sites. Upward and downward arrows indicate increased and decreased enrichment of the indicated proteins in primordial germ cells resulting from exposure of pregnant F0 dams to TBT. **c**, Differential chromatin interaction (DCI) scores of the whole chromatin 19 in mouse primordial germ cells isolated from F1-F3 male or female embryos. DCI scores were determined by Bart3D with 40 kbp bins in chromosome 19 and rolling averaged with 11 bins. Smoothened DCI scores are plotted for the whole chromosome. Blue horizontal bars indicate the location of differential chromatin contacts surrounding the *Ide* gene.**d**, Chromatin contact plots of primordial germ cells isolated from F1-F3 embryos after F0 exposure to TBT (top) or DMSO (bottom). Arrows indicate DCCs gained by the F0 exposure to TBT. **e**, Gene distributions near the transgenerationally conserved DCCs caused by F0 exposure to TBT. The Ide gene is shown in red.

Next, we attempted to identify TAD alterations persistent throughout PGCs isolated from F1-F3 embryos. Applying a strict criterion of differential TAD (dTAD) detection (Differential Chromatin Contact Scores > 3.0), we identified 20 autosomal dTADs conserved between F1 and F2 male PGCs and only one dTAD in chromosome 19 that was conserved across all generations (F1-F3) of male PGCs (Extended data Fig. 3c).

The single dTAD conserved across the F1-F3 PGCs was located in a 10-Mb region of chromosome 19 (30 - 40 Mb) showing significant disturbance in the PGCs of the TBT-exposed group whereas other TADs in this region were strictly conserved across PGCs of the DMSO-exposed control group (Fig. 2c, d). This region contains the *Ide* gene, which encodes insulin degrading enzyme (Fig. 2e). Ide is responsible for much of insulin clearance^16^ and also plays a role in degradation of β-amyloid proteins. Detailed examination of chromatin contacts near the *Ide* gene identified two differential contacts, one of which was conserved across all generations (F1-F3) whereas the other was conserved only in the F1 and F2 generations (Fig. 2d). Formation of significant TAD boundaries near *Ide* was confirmed by visual inspection of the contact diagrams of male PGCs isolated from F1-F3 embryonic testes although the newly formed contact became weaker in F3 embryonic testes (Fig. 2d). These results demonstrated formation of transgenerational, germline-transmitted alterations in chromatin contacts after exposure of pregnant F0 female mice to the obesogen, TBT.

## Ancestral TBT exposure induced TAD formation in the *Ide* gene in male livers

To determine whether the dTADs identified in PGCs were also found in the *Ide* gene in somatic tissues, we sought to examine CTCF binding status in F3 livers because liver is the primary site of *Ide* expression and F2 PGCs give rise to F3 descendants. Five CTCF binding sites predicted by the JASPAR Transcription Factor Binding Site Database lie near the TAD boundary (A-E) together with two known CTCF binding sites (F, G) (Chr19:37,320,000 in mm10) (Fig. 3a). ChIP-qPCR analysis was performed on CTCF or RAD21 chromatin pulldown samples. We observed increased CTCF binding in four regions (C, D, F, G) within the *Ide* gene in male livers and infer that these may form a small chromatin loop (Fig. 3b). No significant enrichment was noted in vehicle group males (Fig. 3b) nor in vehicle or TBT female samples (Fig. 3c). We did not detect enrichment of RAD21 binding on the *Ide* gene (Fig. 3d, e).

**Fig. 3.**
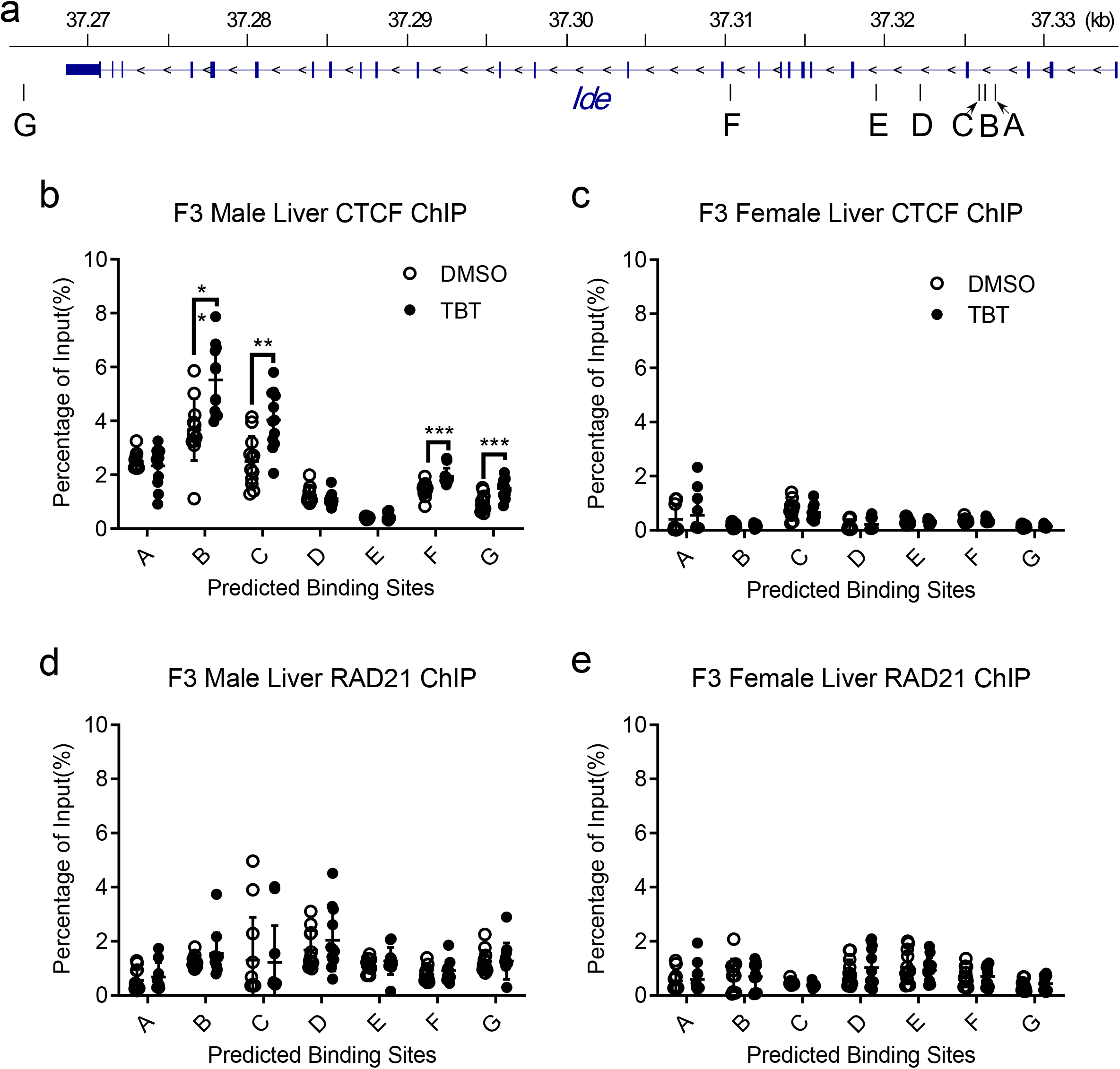
Mice ancestrally exposed to TBT showed increased CTCF binding at the *Ide* gene on chromosome 19. **a**, Five potential CTCF binding sites (A to E) in *Ide* gene of chromosome 19 predicted by JASPAR on the UCSC Genome Browse and two known CTCF binding sites (F and G) were analyzed. Chromatin immunoprecipitation (ChIP) and quantitative real time RT-PCR (qPCR) assays using antibody against CTCF in F3 **b**, male or **c**, female descendants livers. ChIP and qPCR assays using antibody against RAD21 in F3 **d**, male or **e**, female descendants livers. Normal rabbit IgG was used as a non-specific antibody control. Unpaired t-tests were used for qPCR analysis. Data are presented as mean ± s.e.m. *p < 0.05; **p < 0.01; ***p < 0.001.

## Ancestral TBT exposure led to a transgenerational, male-specific hyperinsulinemia

Formation of transgenerationally persistent, novel TAD boundaries in male PGCs within the *Ide* gene (Fig. 2, Extended data Figs. 3, 4) prompted us to hypothesize that expression of *Ide* mRNA might be affected by ancestral TBT exposure. Strikingly, significant decreases in *Ide* mRNA expression were found in the livers of F2 and F3 adult males (Fig. 4a) but not in gonadal WAT (Fig. 4b), skeletal muscle (Fig. 4c) or in females other than F2 female livers (Extended data Fig. 6a-c). Notably, *Ide* mRNA expression was also decreased in F3 male livers from the previously published T3 experiment^12^ (Extended data Fig. 6d).

**Fig. 4.**
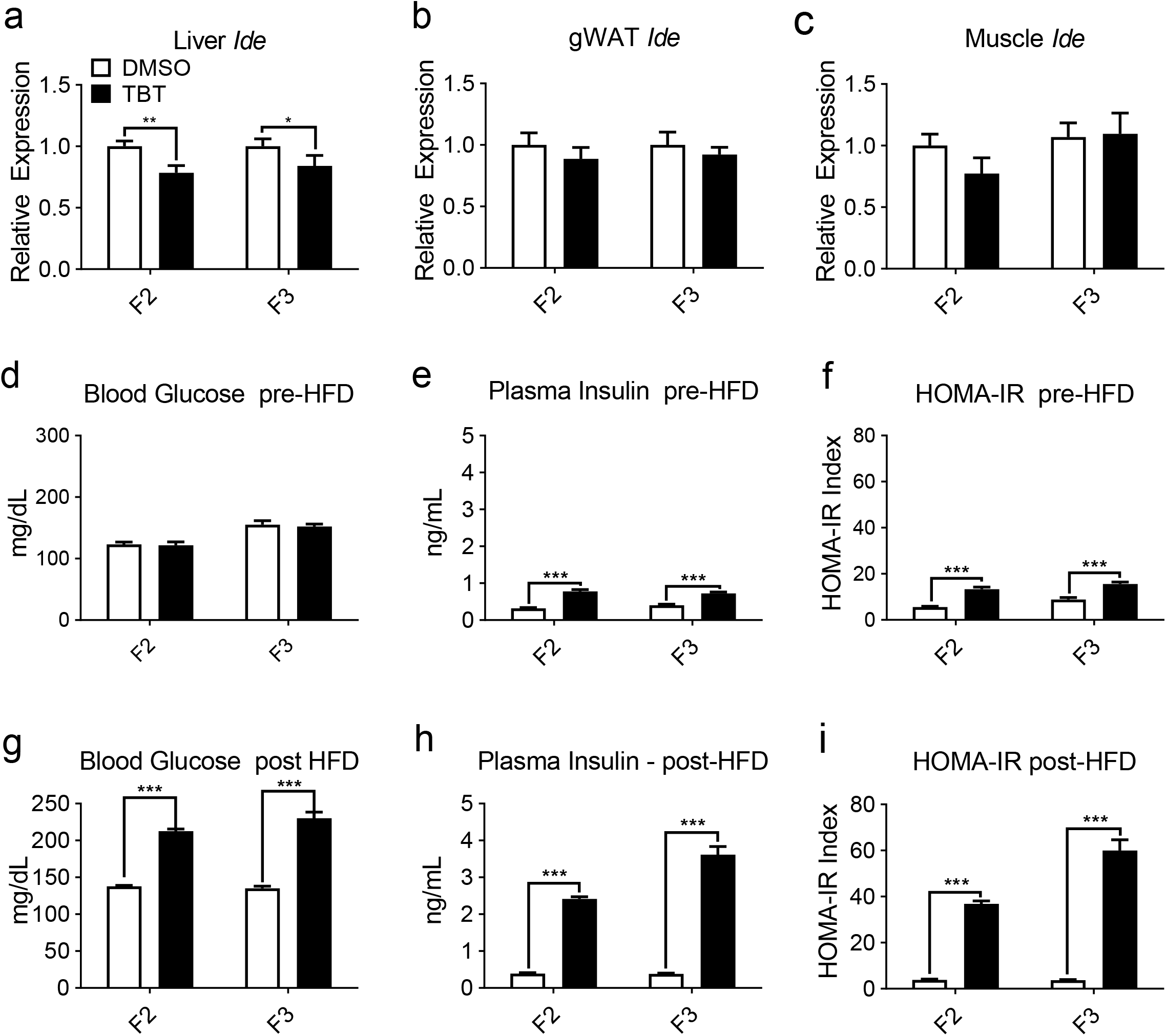
Male mice ancestrally exposed to TBT showed decreased *Ide* gene expression in liver and suffer from hyperinsulinemia before and after six weeks of diet challenge. The relative mRNA levels of *Ide* gene were assayed by quantitative PCR in **a**, liver, **b**, gonadal white adipose tissue (gWAT), and **c**, soleus muscle of F2 (n = 15) and F3 (n =16) male descendants with the expression normalized to GAPDH. Data are expressed as mean fold change ± SEM and assayed in duplicate. *p < 0.05 and **p < 0.01, compared with DMSO groups by unpaired t-test. **d**, Plasma glucose and **e**, insulin of male mice before diet challenge (F2 at 5 weeks age and F3 at 17 weeks age) were measured and **f**, HOMA-IR indexes were calculated accordingly. **g**, Plasma glucose and **h**, insulin of male mice after six weeks of diet challenge (F2 at 11 weeks age and F3 at 22 weeks age) were measured and **i**, HOMA-IR indexes were calculated accordingly. Statistical significance was determined using two-way ANOVA. Pair-wise Bonferroni post-tests were used to compare different groups. Data are presented as mean ± s.e.m. *p < 0.05; **p < 0.01; ***p < 0.001.

No differences in fasting blood glucose levels were observed in either males (Fig. 4d) or females (Extended data Fig. 7a) prior to diet challenge, as we previously reported^8^. However, plasma insulin levels in TBT group males were significantly higher (Fig. 4e), in accord with the decreased hepatic *Ide* mRNA expression (Fig. 4a) whereas no significant differences were observed in females (Extended data Fig. 7b). We calculated homeostasis model assessment of insulin resistance (HOMA-IR) to estimate insulin resistance and noticed that TBT-group male descendants showed higher potential for insulin resistance than did DMSO descendants (Fig. 4f). Females showed no differences (Extended data Fig. 7c). After 6 weeks of HFD diet, TBT male descendants were hyperglycemic (Fig. 4g) and hyperinsulinemic (Fig. 4h), in addition to having increased fat content (Fig. 1a, b); females showed no effects (Extended data Fig. 7 d, e). HOMA-IR index was strikingly increased in the male (Fig. 4i) but not female (Extended data Fig. 7f) TBT animals, suggesting a high likelihood of insulin resistance compared with vehicle controls.

To address whether insulin secretion or insulin breakdown was responsible for increased insulin levels in TBT-group males, we measured C-peptide (Extended data Fig. 8) and calculated C-peptide:insulin ratios (Extended data Fig. 9). Analysis of C-peptide levels revealed that there were no differences between DMSO and TBT group males or females before HFD challenge, or females after HFD challenge (Extended data Fig. 8). C-peptide levels increased ∼50% in DMSO and TBT-group males after HFD challenge, indicating that HFD led to increased insulin secretion. Measurement of the ratios of C-peptide:insulin indicated that increased insulin levels observed in TBT-group males before and after HFD were likely the result of impaired insulin clearance resulting from decreased IDE production (Extended data Fig. 9a, b). In accord with our previous publications^8,12^, plasma leptin levels were strongly increased in TBT-group males after HFD challenge (Extended data Fig. 10).

## Discussion

Obesogens are chemicals that lead to increased WAT mass in exposed organisms^2,17^. TBT activated the nuclear receptors peroxisome proliferator activated receptor gamma (PPARγ) and its heterodimeric partner, retinoid ‘X’ receptor (RXR)^18,19^, leading to increased commitment of multipotent mesenchymal stromal stem cells to the adipose lineage and differentiation of pre-adipocytes into mature adipocytes^20^. TBT promoted fat accumulation in vivo in a variety of model systems, including mice^2,17^. Our previous experiments demonstrated increased WAT accumulation when dietary fat was elevated modestly (21.6% vs 13.1% calories from fat) in male F3 and F4 descendants of pregnant F0 mouse dams exposed to TBT throughout pregnancy^11,12^ or pregnancy and lactation^8,10,13^. TBT exposure of pregnant F0 dams resulted in a stable, male-specific predisposition to obesity in exposed (F1 were exposed in utero, F2 were exposed as germ cells in F1) and unexposed (F3, F4) descendants.

We previously identified blocks of iso-directional, differentially methylated DNA (isoDMBs) in WAT of F4-generation male mice after exposure of F0 dams to TBT throughout pregnancy and lactation^8,10^. Genomic DNA regions in WAT where isoDMBs were under-methylated compared to controls were enriched in metabolic genes such as leptin and these regions were less accessible in F3 and F4 sperm of the TBT group than in controls^8^. We proposed that ancestral TBT exposure caused changes in higher-order chromatin structure that were then inherited or reconstructed every generation, ultimately resulting in changes in chromatin accessibility and DNA methylation that altered expression of adipogenic and metabolic genes compared with controls^8,10^.

Transgenerational inheritance or reconstruction of altered higher-order chromatin structure offers an attractive unifying model that provides a common basis to integrate how disparate mechanisms such as DNA methylation, histone methylation, histone retention, and noncoding-RNA expression might be coordinately regulated across the generations. Higher-order chromatin structure is reflected in the presence or absence of chromatin TADs and loops that modulate accessibility to DNA and histone-modifying enzymes, to histones and to the transcription machinery^15^. Support for such a model required the identification of persistently altered regions of higher-order chromatin structure. Here we used genome-wide Hi-C and BART-3D analysis to identify TADs whose presence was stably altered in PGCs by direct or ancestral TBT exposure. Critically, the most high-scoring TAD identified objectively by BART-3D analysis was on chromosome 19, within the *Ide* gene. While we have only assessed the presence of TADs in PGCs from the current (T4) experiment, expression of *Ide* was also reduced in livers from a previous experiment (T3)^12^. Reduced *Ide* expression did not affect fasting glucose levels in the current T4 experiment, but significantly altered basal insulin levels, then led to strong increases in both glucose and insulin levels in HFD-challenged F2 and F3 males. This increase in insulin was not the result of increased insulin secretion because C-peptide levels did not change between treatment groups. Since only the male animals in our transgenerational experiments responded to diet challenge and then only increased WAT mass after HFD diet was initiated, decreased hepatic *Ide* expression in males appears to be a strong component of the transgenerational susceptibility to obesity. Previous gene knockout studies confirmed a role for IDE in insulin clearance; *Ide* loss-of-function produced hyperinsulinemia and age-dependent glucose-intolerance^16^. It is also notable that leptin levels were increased in TBT-group males since it is known that hyperinsulinemia and insulin resistance impair leptin signaling, leading to leptin resistance^21^.

Together with our previous studies^8,10^, the new data presented here support a model in which transgenerational non-genetic propagation of environmentally-induced phenotypes relied on alterations in chromatin structure. Altered chromatin structure necessarily changes the accessibility of DNA and histones to modifying enzymes such as DNA and histone methyl transferases, the location and retention of histones and the expression of various genes, including those for non-coding RNAs. These mechanisms could interact and be preserved across generations and in various types of differentiated cells^10^.

## Methods

### Chemicals and Reagents

TBT, dexamethasone, isobutylmethylxanthine, insulin were purchased from Sigma-Aldrich (St. Louis, MO). Rosiglitazone (ROSI) was purchased from Cayman Chemical (Ann Arbor, MI). Embryoid Body Dissociation Kit (#130-096-348) was purchased from Miltenyi Biotec (North Rhine-Westphalia, Germany). Zombie Red Dye (#77475) was purchased from BioLegend (San Diego, CA). PE Mouse anti-SSEA-1 (#560142) and Alexa-Fluor 647 Mouse anti-CD61 (#563523) were purchased from BD Biosciences (Franklin Lakes, NJ). Blood glucose meter kits (BG1000) were purchased from Clarity Diagnostics (Boca Raton, FL). Mouse Leptin ELISA Kit (#90030), Mouse C-peptide ELISA kit (#80954) and mouse insulin ELISA kit were purchased from Crystal Chem (Elk Grove Village, IL, USA). Arima-HiC Kit (A510008) was purchased from Arima Genomics (San Diego, CA). Ultra-pure formaldehyde (#18508) was purchased from Ted Pella Inc (Redding, CA). MinElute Reaction Cleanup Kit (#28206) was purchased from Qiagen (Hilden, Germany).

### Animal maintenance and exposure

C57BL/6J mice were purchased from the Jackson Laboratory (Sacramento, CA) and housed in micro-isolator cages in a temperature-controlled room (21–22 °C) with a 12 h light/dark cycle. Water and food were provided *ad libitum* unless otherwise indicated. Animals were treated humanely and with regard for alleviation of suffering. All procedures conducted in this study were approved by the Institutional Animal Care and Use Committee of the University of California, Irvine. At the moment of euthanasia, each mouse was assigned a code, known only to a lab member not involved in the dissection process. All tissue harvesting was performed with the dissector blinded to which groups the animals belonged. Group sizes were based on our prior experiments and a priori power analysis

For this new transgenerational experiment, denoted as T4; we purchased 50 male and 148 female C57BL/6J mice (5 weeks of age). Female mice (74 females per treatment group) were randomly assigned to the different F0 treatment groups and exposed via drinking water to 50 nM TBT or 0.1% DMSO vehicle (both diluted in 0.5% carboxymethyl cellulose in water to maximize solubility), for 7 days prior to mating as we have described^8,12^. One male was housed with two vehicle or 50 nM TBT exposed F0 females per cage to breed during the dark cycle (6PM to 6AM). Vaginal plug appearance was defined as embryonic day (E) 0.5. Treatment was removed during mating, then resumed for F0 females after copulation plugs detected (and males removed) then maintained until pups were born (Extended data Fig. 11). This TBT concentration was chosen based on our previous studies^8,11,12,22^ and is five-fold lower than the established no observed adverse effect level (NOAEL)^23^. While chemicals were administered to the dams throughout pregnancy, sires were never exposed to the treatment. No statistically significant differences were observed in the number of pups or the sex ratio per litter among the different groups (Extended data Fig. 12). It should be noted that F2 descendants were exposed to TBT as germ cells in the exposed F1 embryos. F3 descendants were not exposed to TBT.

From each generation, we randomly chose only 1 male and 1 female per litter for endpoint analysis and another 1 male and 1 or 2 females per litter for breeding to produce the next generation. There were insufficient animals in the F1 generation to both breed the F2 generation and analyze phenotypes in the diet challenge, so we only bred the F1 animals. To randomize the breeding process as much as possible, we did not breed siblings and never bred females from the same litter with the same male. Control animals were bred to each other and TBT-exposed animals were bred to each other.

### Diet challenge and body composition analysis

Animals from control and treatment groups were maintained on a standard diet (SD) (PicoLab 5053; 24.5% Kcal from protein, 13.1% Kcal from fat, and 62.3% Kcal from carbohydrates) from weaning onward. In diet challenge experiments, F2 (14 males and 14 females for each group) and F3 (15 males and 15 females for each group) were switched to higher fat diet (HFD) (PicoLab Rodent Chow, 5058) whereas control groups (F2: 15 males and 15 females; F3: 12 males and 12 females) were maintained on the SD (PicoLab Rodent Chow, 5053). Body weight and body composition were measured weekly for each animal using EchoMRI™ Whole Body Composition Analyzer, which provides lean, fat and water content information. Total water weight includes free water mainly from the bladder and water contained in lean tissue. F2 descendants started diet challenge at 4 weeks of age for 8 weeks when a significant fat content increase was confirmed and persisted. F3 descendants starting the diet challenge at week 5 had not become significantly fatter by week 17. Therefore, we switched the control F3 animals to the HFD at week 17 for 5 weeks. Fat content was significantly increased in this group by week 19. Mice were fasted for 12 hours prior to euthanasia and tissue collection.

Blood was collected via the saphenous vein at week 4 and week 12 (before and after diet challenge) for F2, and at weeks 4, 12, and 22 for F3. Blood was collected into heparinized tubes, then centrifuged for 15 minutes at 5,000 RPM at 4ºC. Resulting plasma was transferred to a clean tube and preserved at -80ºC. Animals were euthanized by isofluorane exposure followed by cardiac exsanguination after 4 hours fasting. Blood was drawn into a heparinized syringe and centrifuged for 15 minutes at 5,000 rpm at 4ºC. Resulting plasma was transferred to a clean tube and preserved at -80ºC. We measured plasma leptin levels to confirm the previously reported phenotypes^8^. Inguinal white adipose tissue (iWAT), gWAT, pancreas, spleen, liver, interscapular brown adipose tissue (iBAT), and soleus muscle were flash frozen in liquid N_2_ then stored at -80ºC for subsequent analysis. Feces was freshly collected from animals prior to, and during the diet challenge at week 4 and week 12 for F2, week 4, 12, and 22 for F3, and stored at - 80ºC.

### PGC isolation

A randomly-selected subset of pregnant females was euthanized 13 days (E13.5) after vaginal plug detection. E13.5 embryos were isolated from euthanized pregnant dams, and E13.5 gonads containing primordial germ cells (PGCs) were isolated using Leica MZ9.5 Binocular Stereo Microscope. Gonads were identified and sexed by their characteristic morphology at E13.5 (Extended data Fig. 13a, b) and sex was verified by PCR^24^. Primer sequences are given in Extended data Table 1. Gonads from same-gender embryos in each litter were pooled prior to tissue dissociation. Gonads were enzymatically digested using Embryoid Body Dissociation Kit (Miltenyi Biotec). Next, total dissociated gonad cells were stained with Zombie Red Dye (BioLegend), PE Mouse anti-SSEA-1 (BD Pharmingen), and Alexa-Fluor 647 Mouse anti-CD61 (BD Pharmingen). Primordial germ cells were purified based on the expression of Zombie Red^−^/SSEA-1^+^/CD61^+^ using BD FACS Aria II Cell Sorter (BD Bioscience). Somatic gonad cells were purified based on the expression of Zombie Red^−^/SSEA-1^−^/CD61. The gating strategy and purity of the isolated cells is shown in Supplementary Extended data Fig. 14.

### Hi-C data generation

Five litters of each group with more than 10,000 PGCs (Red^−^ /SSEA-1^+^/CD61^+^) were designated to proceed for Hi-C sample preparation using Arima Hi-C Kit following the manufacturer’s instructions for low-input Hi-C sequencing. This low-input protocol supported quantitative determination of the topologically associating domains (TADs) from 10,000 human cells (Arima Genomics Application Note: “Unlock Low-Input 3D Genome Analysis with the Arima-HiC Kit”, Arima Genomics), which was confirmed by our current study (Extended data Fig. 2). Briefly, cells were fixed with formaldehyde (Ted Pella) to crosslink chromatin contacts. Genomic DNA was isolated from the fixed cells and digested using a restriction enzyme cocktail. The 5’-overhangdigested ends were filled with biotinylated nucleotides, and spatially proximal digested ends were ligated. Proximally ligated DNA fragments, which capture chromatin contacts were purified, fragmented by sonication, and enriched using streptavidin-conjugated beads. Illumina sequencing libraries were synthesized from the solid phase-captured DNA fragments using the Swift Biosciences® Accel-NGS® 2S Plus DNA Library Kit (Swift). Libraries were sequenced using an Illumina NovoSeq 6000 deep sequencer to obtain 150 + 150 nt paired-end FASTQ reads.

### Identification of differential topologically associating domains (dTADs) and genes

**-** After adaptor sequences and low-quality reads (<30) were removed using the Trim Galore! tool, FASTQ reads were subjected to Hi-C seq analysis using the Hi-C Pro tool^25^. The Hi-C Pro quality control plots showed that least 60 million valid pairs were generated for each library with greater than 60% long (>20 kb) *cis* interactions, indicating successful generation of sufficient amounts of high-quality Hi-C data (Extended data Figs. 2e-g). Using these data, we were able to reproduce a known TAD profile surrounding the *HoxD* gene cluster^14^, which was demonstrated using a contact map (Extended data Fig.2h) and a directionality index plot (Extended data Fig.2i). Locations of chromatin contact boundaries were predicted using the clusterTAD tool^26^, which accurately predicts TADs in mammalian cells using an unsupervised machine learning algorithm. Enrichment of DNA-associating proteins at the differential TAD boundaries between TBT versus vehicle groups was predicted using BART-3D^27^. Distributions of TADs were visualized using the OmicCircos^28^ and the Sushi^29^ R/Bioconductor tools.

### Chromatin immunoprecipitation-Quantitative Polymerase Chain Reaction

Chromatin immunoprecipitation (ChIP) was performed using the method by Abcam and optimized with the established method we previously described^30^. Briefly, 50 mg of liver tissue that had been snap-frozen in liquid N_2_ were thawed on ice in cold PBS and dispersed into single cell suspensions using a 100 μm cell strainer (#22363549; Fisher Brand, PA). Cells were washed twice with PBS containing protease inhibitor cocktail (#ab201111; Abcam, Cambridge, UK) then resuspended and fixed at room temperature for 10 minutes with 1% paraformaldehyde (Fisher Chemical, PA) in DMEM, followed by an ice-cold phosphate-buffered saline wash, and then quenched for 5 minutes with 125 mM glycine at room temperature. Fixed cells were washed, collected by centrifugation, then resuspended in phosphate-buffered saline at 10^7^ cells/mL. To isolate nuclei, cell pellets were lysed at 4°C for 10 minutes with a gentle detergent recipe consisting of 50 mM HEPES-KOH, pH 7.5, 140 mM NaCl, 1 mM EDTA, 10% glycerol, 0.5% Nonidet P-40, 0.25% Triton X-100, Protease Inhibitor Cocktail (#ab201111; Abcam, Cambridge, UK). Nuclei were recovered by centrifugation at 8000 x g for 15 minutes, washed for 10 minutes at room temperature (10 mM Tris-HCl, pH 8.0, 200 mM NaCl, 1 mM EDTA, 0.5 mM EGTA, protease inhibitors (#ab201111; Abcam, Cambridge, UK), and lysed in 300 μL nuclear lysis buffer (10 mM Tris-HCl, pH 8.0, 200 mM NaCl, 1 mM EDTA, 0.5 mM EGTA, 0.1% Na-deoxycholate, 0.5% N-lauroylsarcosine, protease inhibitors (#ab201111; Abcam, Cambridge, UK). Chromatin samples were prepared by sonicating in 0.5 mL thin-walled polymerase chain reaction tubes (BrandTech, CT) using a QSonica Q800R2 (QSonica, CT) with the following settings: 30 seconds on/30 seconds off, amplitude 40% repeated for 30 minutes. Triton X-100 (1%) was added to sonicated lysates prior to high-speed, cold centrifugation to remove debris. A total of 5 μg DNA was immunoprecipitated with preblocked protein A/G Dynabeads (Thermo Fisher Scientific, MA) complexed to 2.5 μg antibody (anti-CTCF, ab128873, anti-RAD21, ab217678, or Isotype IgG control, ab171870, Abcam, Cambridge, UK). Beads were washed three times with LiCl buffer (50 mM HEPES-KOH, pH 7.5, 500 mM LiCl, 1 mM EDTA, 1% Nonidet P-40, 0.7% Na-deoxycholate) and once with Tris-EDTA buffer plus 50mM NaCl. To release chromatin from beads, pelleted beads were resuspended in elution buffer (50mM Tris-HCl, pH 8.0, 10 mM EDTA, 1% sodium dodecyl sulfate) and incubated at 65°C for 30 minutes. Cross-link reversal was performed overnight at 65°C. DNA samples were purified using Qiaquick PCR Cleanup kit (#28106, Qiagen, Germantown, MD) following RNase A (0.2 mg/mL, 2 hours, 37°C) and proteinase K (0.2 mg/mL, 2 hours, 55°C) treatment. Input DNA content was determined by spectrophotometry (Nanodrop, Thermo Fisher Scientific, MA). For analysis of candidate loci, real-time PCR was performed using SYBRTM Green PCR Master Mix (Thermo Fisher Scientific, MA) on a Roche LightCycler 480 II (Roche, Switzerland) according to the recommended protocol. Enrichment of the ChIP target were presented as fold difference between specific Ab-immunoprecipitated samples and the immunoprecipitated total input with an IgG control. Primer sequences of the examined loci are listed in Extended data Table 1. Multiple primer sets were tested for each site. For sites B and C, 2 of 2 primer sets showed significant enrichment. For binding site F, 1 of 3 primer sets showed enrichment and for site G, 2 of 3 showed enrichment.

### Quantitative real time reverse transcriptase polymerase chain reaction

Tissue that had been previously snap-frozen in liquid N_2_ was cut into ∼20 mg pieces and lysed with Trizol following the manufacturer’s recommended protocol (Thermo Fisher Scientific, MA); total RNA was recovered after isopropanol precipitation (Fisher Chemical, PA). Complementary DNA was synthesized from 5 μg total RNA using SuperScript IV First-Strand Synthesis System (Thermo Fisher Scientific, MA) according to the manufacturer’s instructions. Gene expression was assessed with real-time quantitative polymerase chain reaction (qPCR) using SYBRTM Green PCR Master Mix (Thermo Fisher Scientific, MA) on a Roche LightCycler 480 II (Roche, Switzerland). Primer sequences of the examined genes were listed in Extended data Table 1. Cycle threshold values were quantified as the second derivative maximum using LightCycler software (Roche, Switzerland). The 2^−ΔΔCt^ method^31^ was used to analyze RT-qPCR data and determine relative quantification corrected for primer efficiency. *Ide* expression was normalized to the housekeeping gene, *GAPDH*, and compared to DMSO descendants group. Error bars represent the SEM from 15 to 17 biological replicates, calculated using standard propagation of error.

### Measurement of insulin and C-peptide

C-peptide serum levels were measured by EIA (Crystal Chem #80954; Elk Grove Village, IL, USA) at two different time points (before and after diet challenge) in plasma from blood samples drawn after overnight (12 hours) fasting. Insulin levels were measured by EIA (Crystal Chem #90080; Elk Grove Village, IL, USA) in plasma from blood samples drawn after overnight (12 hours) fasting.

## Acknowledgements

Correspondence and requests for materials should be addressed to B.B. or T.S. Supported by grants from the National Institutes of Health, USA (R01ES023316 and R01ES031139) to B.B. and T.S. We thank Drs. Sha Sun (University of California, Irvine, CA USA), Xiao-Min Ren (Kunming University of Science and Technology, China), Angel Nadal (Universidad Miguel Hernández de Elche. Spain), Ivan Quesada (Universidad Miguel Hernández de Elche. Spain), Hanjun Lee (Whitehead Institute, Cambridge, MA, USA), Michael Lawrence (Whitehead Institute, Cambridge, MA, USA) and Rob Lustig (University of California, San Francisco, CA USA) for critically reading the manuscript and Dr. Hanjun Lee for suggesting BART-3D. We wish to thank Drs. Raquel Chamorro-Garcia (University of California, Santa Cruz) and Grant MacGregor (University of California, Irvine) for advice and assistance during the early stages of this project.

## Author contribution

R.C.C., R.J.E., T.S. and B.B. designed the experiments reported in this manuscript. R.C.C., R.J.E., E.J., Y.H., H.B.W., A.N., K.S., J.O., T.S., and B.B. performed experiments. R.C.C., R.J.E., T.S. and B.B. analyzed data. R.C.C., T.S. and B.B. wrote the manuscript.

## Data availability statement

Hi-C datasets will be available at Gene Expression Omnibus (xxxxxx).

## Figure legends

**Extended data Fig. 1.**
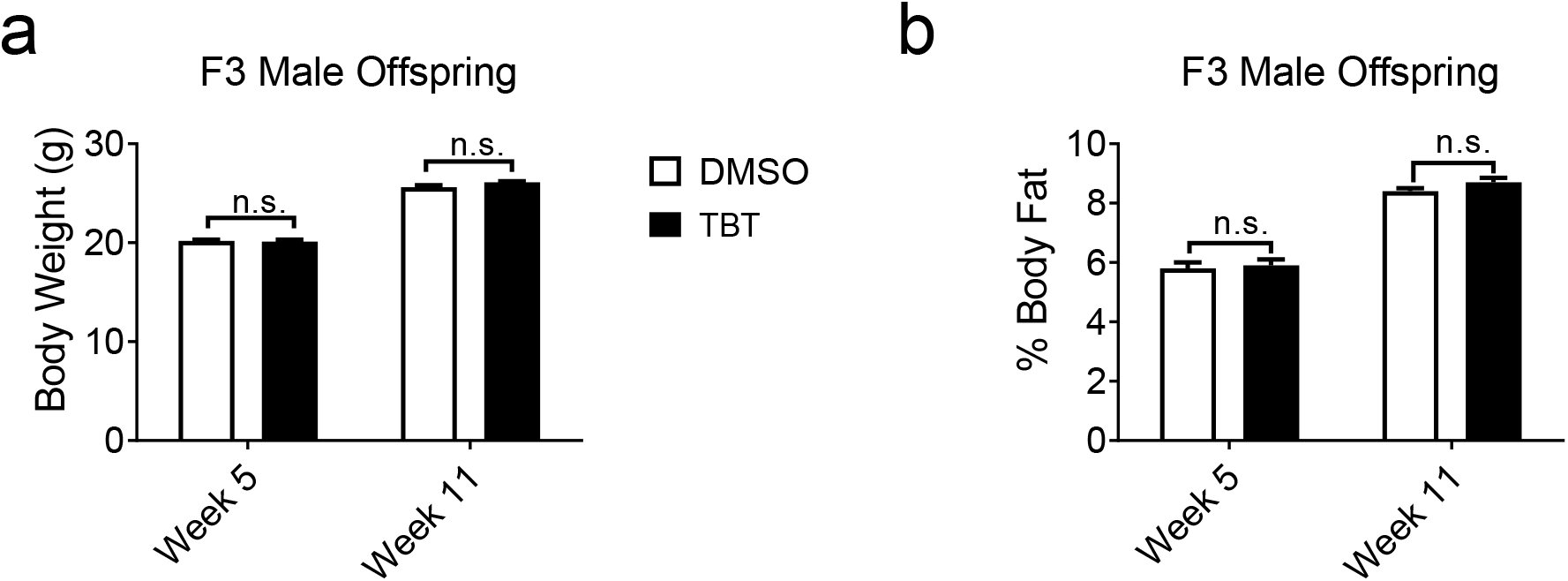
Mice ancestrally exposed to TBT showed no effects on body weight or fat content under diet challenge during early ages of F3. **a**, Body weight and **b**, relative body composition of F3 male descendants (n = 16) at 5 weeks age (before diet challenge) and 11 weeks age (after six weeks of diet challenge) were measured. Statistical significance was determined using two-way ANOVA. Pair-wise Bonferroni post-tests were used to compare different groups. Data are presented as mean ± s.e.m.

**Extended data Fig. 2.**
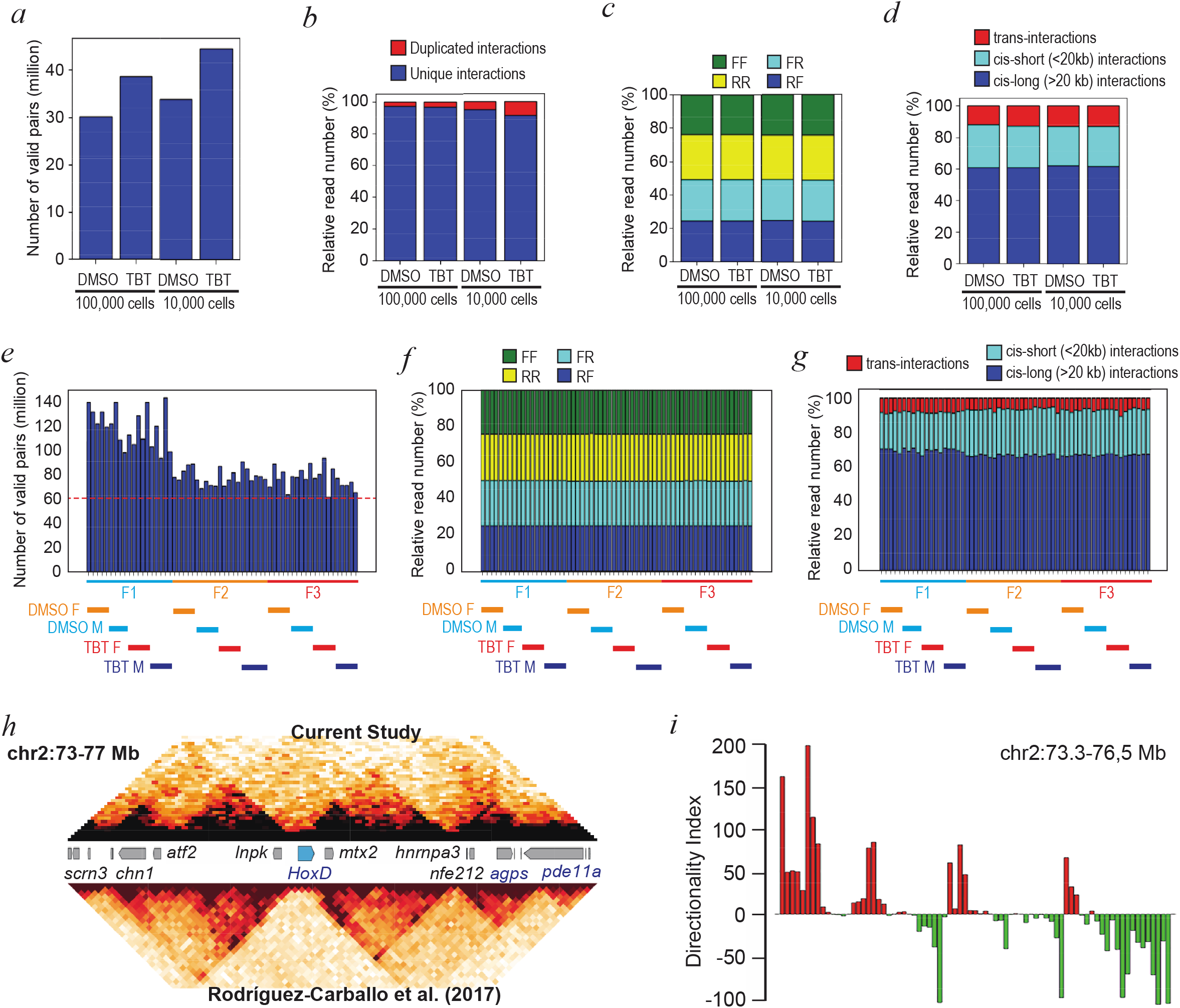
Quality assessments of Hi-C data generated from E13.5 mouse primordial germ cells. **a-d**, Hi-C Pro quality measure outputs for Hi-C data of mouse primordial germ cells isolated from F1 embryos after gestational exposure of F0 mothers to DMSO or TBT. Primordial germ cells were pooled from multiple E13.5 F1 embryos and enriched by FACS to obtain 100,000 or 10,000 cells. **a**, Numbers of valid pairs of reads. **b**, Proportions of duplicated or unique chromatin interactions. **c**, Strands sequenced in reads representing unique interactions: F, forward; R, reverse. An unbiased Hi-C experiment involves equal proportions of the four F x R combinations of strand sequencing. **d**, Proportions of unique chromatin interactions. Greater than 60% of reads represented long (>20 kb) cis interactions, which correspond to the size of loops (< 0.2 Mb) or Topologically Associating Domains (TADs, 0.2 – 1 Mb). **e-g**, Hi-C Pro quality measure outputs for Hi-C data generated from ∼20,000 FACS-enriched E13.5 mouse primordial germ cells isolated from F1, F2, and F3 male (M) and female (F) embryos after gestational exposure of F0 mothers to DMSO or TBT. **e**, Numbers of valid pairs of reads. At least 60 million valid pairs of reads were generated for each embryo. **f**, Strands sequenced in reads representing unique interactions. **g**, Proportions of unique chromatin interactions. **h**,**i**, Detection of known TADs surrounding the mouse HoxD gene cluster. **h**, Chromatin interaction plots of the current study (top; DMSO-group F2 male) and a published study (bottom). **i**, Directionality indexes surrounding the HoxD gene cluster calculated from data of the current study.

**Extended data Fig. 3.**
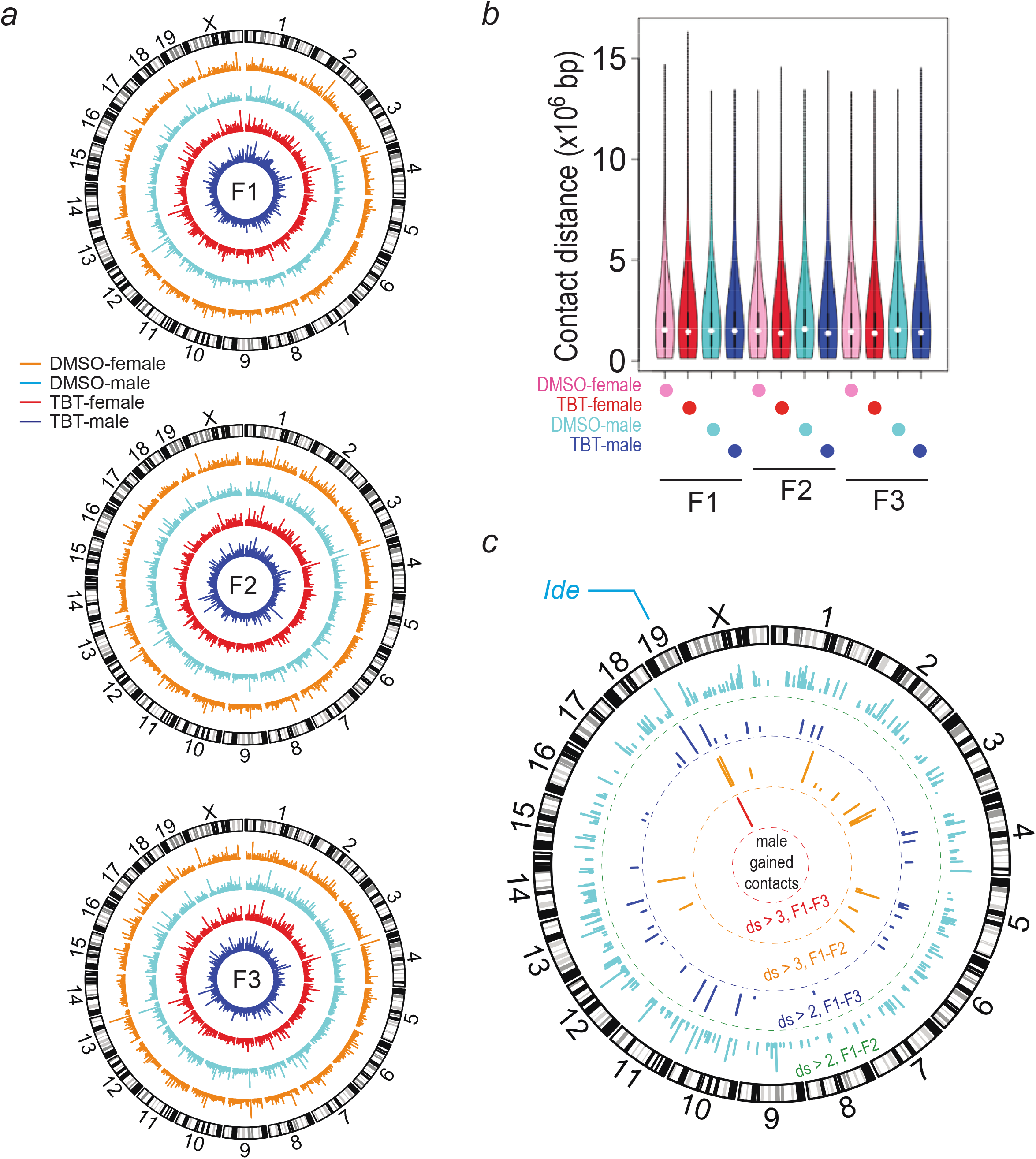
Global distribution of chromatin contacts in the genome of mouse primordial germ cells. **a**, Circos plots of chromatin contact locations and distances in primordial germ cells isolated from F1-F3 embryos. Length of bars indicates distances between chromatin contacts at each chromosomal location. **b**, Violin plots of distances between chromatin contacts. **c**, Circos plot of DCCs gained in the genome of male primordial germ cells by F0 exposure to TBT. Length of each bar indicates differential scores (ds). Concentric arches from the periphery to the center show DCCs with relaxed to strict criteria – namely, ds>2 conserved between F1-F2 (cyan) or F1-F3 (blue), ds>3 conserved between F1-F2 (orange) or F1-F3 (red). Chromosomal locations of the *Ide* gene encoding the insulin degrading enzyme is indicated outside the circular ideograms.

**Extended data Fig. 4.**
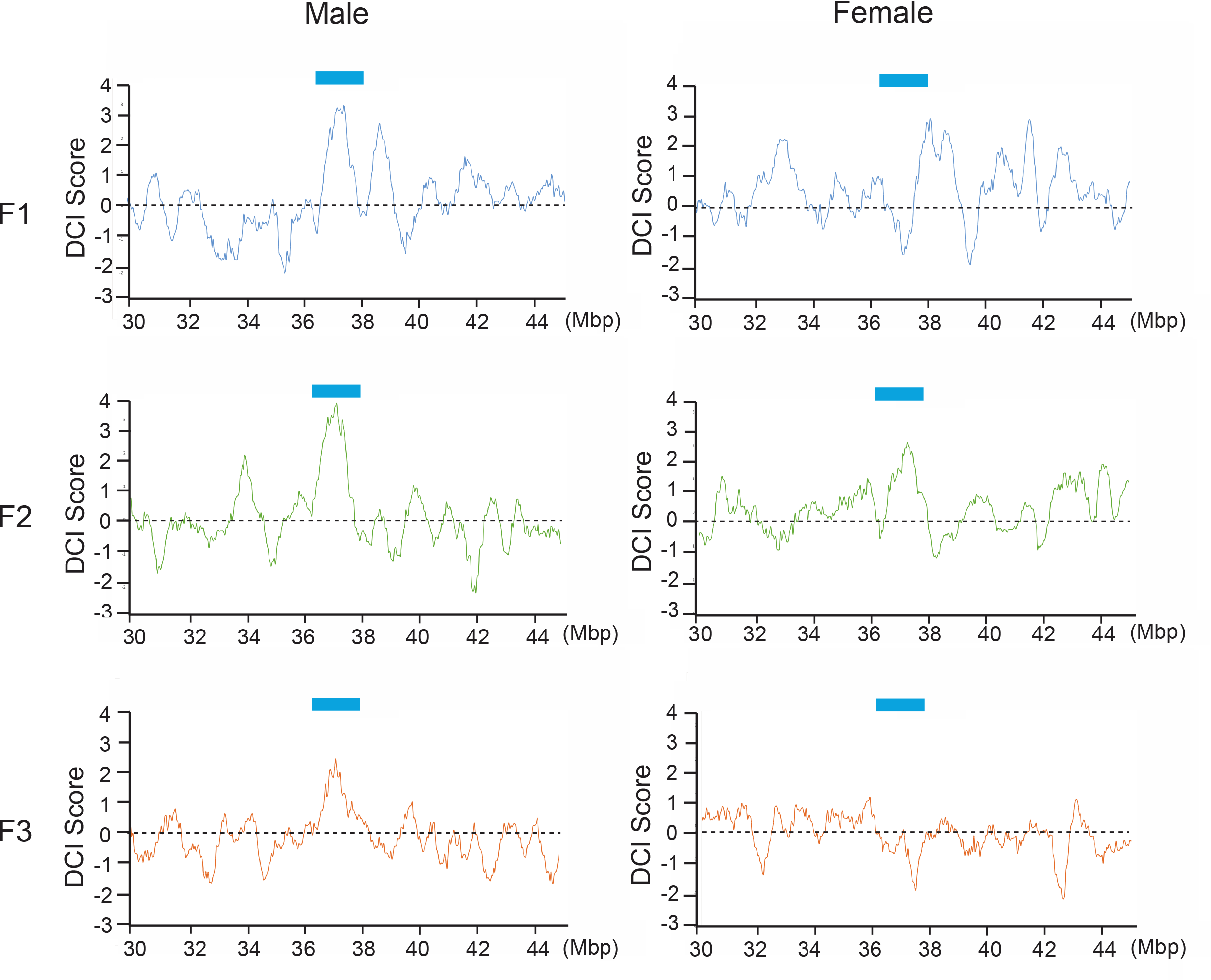
Differential chromatin interaction (DCI) scores of a part of chromatin 19 surrounding the *Ide* gene in mouse primordial germ cells isolated from F1-F3 male or female embryos. DCI sores were determined by Bart3D with 40 kbp bins in chromosome 19 and rolling averaged with 11 bins. Smoothened DCI scores are plotted for chr19: 30-44.5 Mbp. Blue horizontal bars indicate the location of differential chromatin contacts surrounding the *Ide* gene.

**Extended data Fig. 5.**
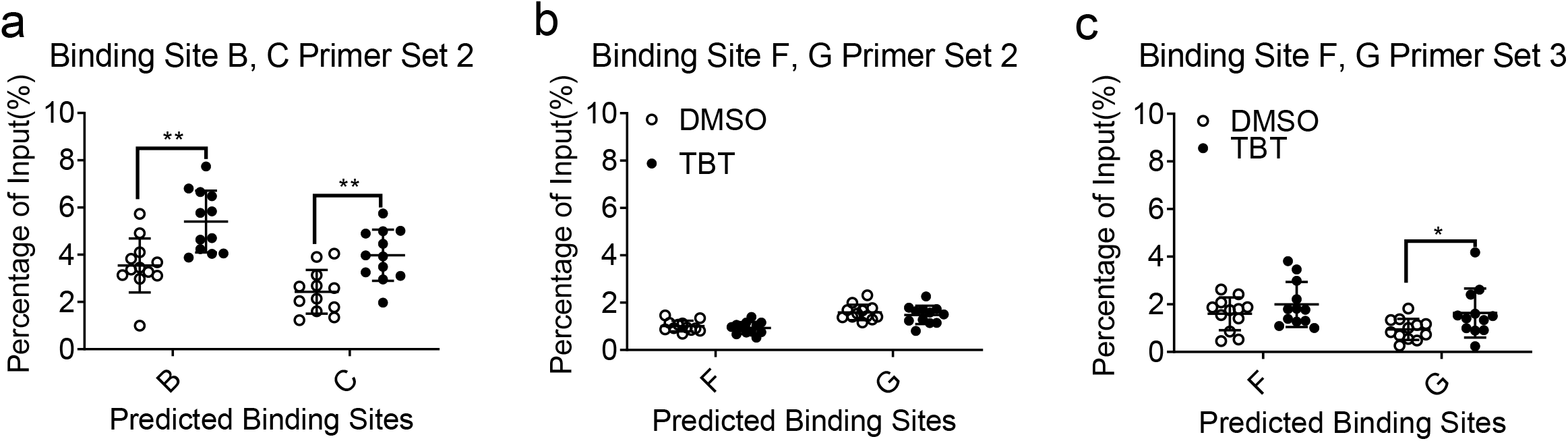
Reproducibility of ChIP qPCR on CTCF binding sites. **a**, The second set of primers was used to test CTCF binding sites B and C. **b**, The second and **c**, the third sets of primers were used to test CTCF binding sites F and G. Unpaired ttests were used for qPCR analysis. Data are presented as mean ± s.e.m. *p < 0.05; **p < 0.01

**Extended data Fig. 6.**
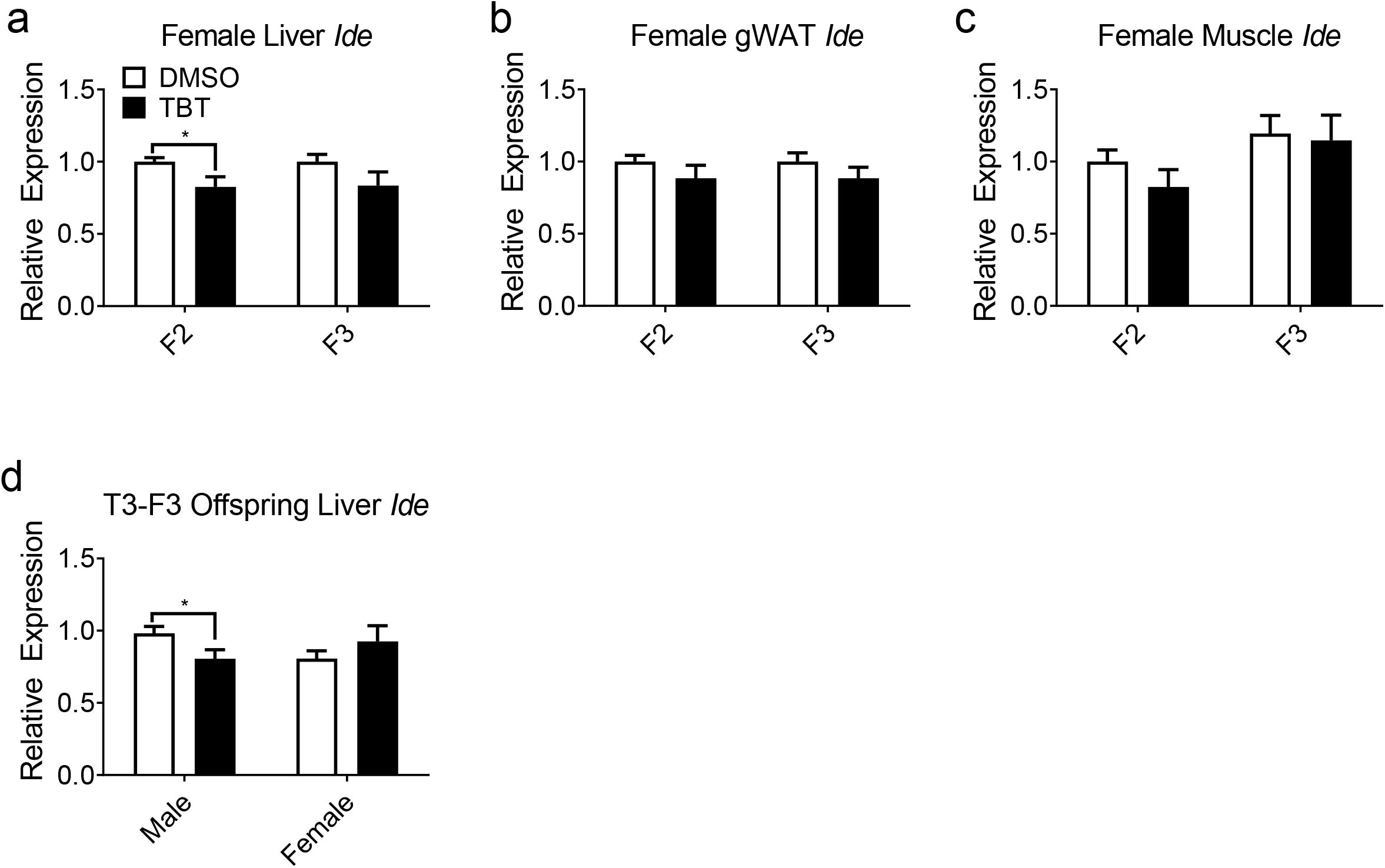
Expression of *Ide* in female tissues and in a previous transgenerational experiment. The relative mRNA levels of the *Ide* gene were assayed by quantitative PCR in **a**, liver, **b**, gonadal white adipose tissue (gWAT), **c**, soleus muscle of F2 (n = 17) and F3 (n =16) female descendants from the current experiment. Panel **d**, shows expression in livers of F3 descendants from the previous third transgenerational study^12^ (DMSO male = 16, TBT male = 16, DMSO female = 14, and TBT female = 11) with the expression normalized to GAPDH. Data are expressed as mean fold change ± s.e.m. and assayed in duplicate. *p < 0.05 compared with DMSO groups by unpaired t-test.

**Extended data Fig. 7.**
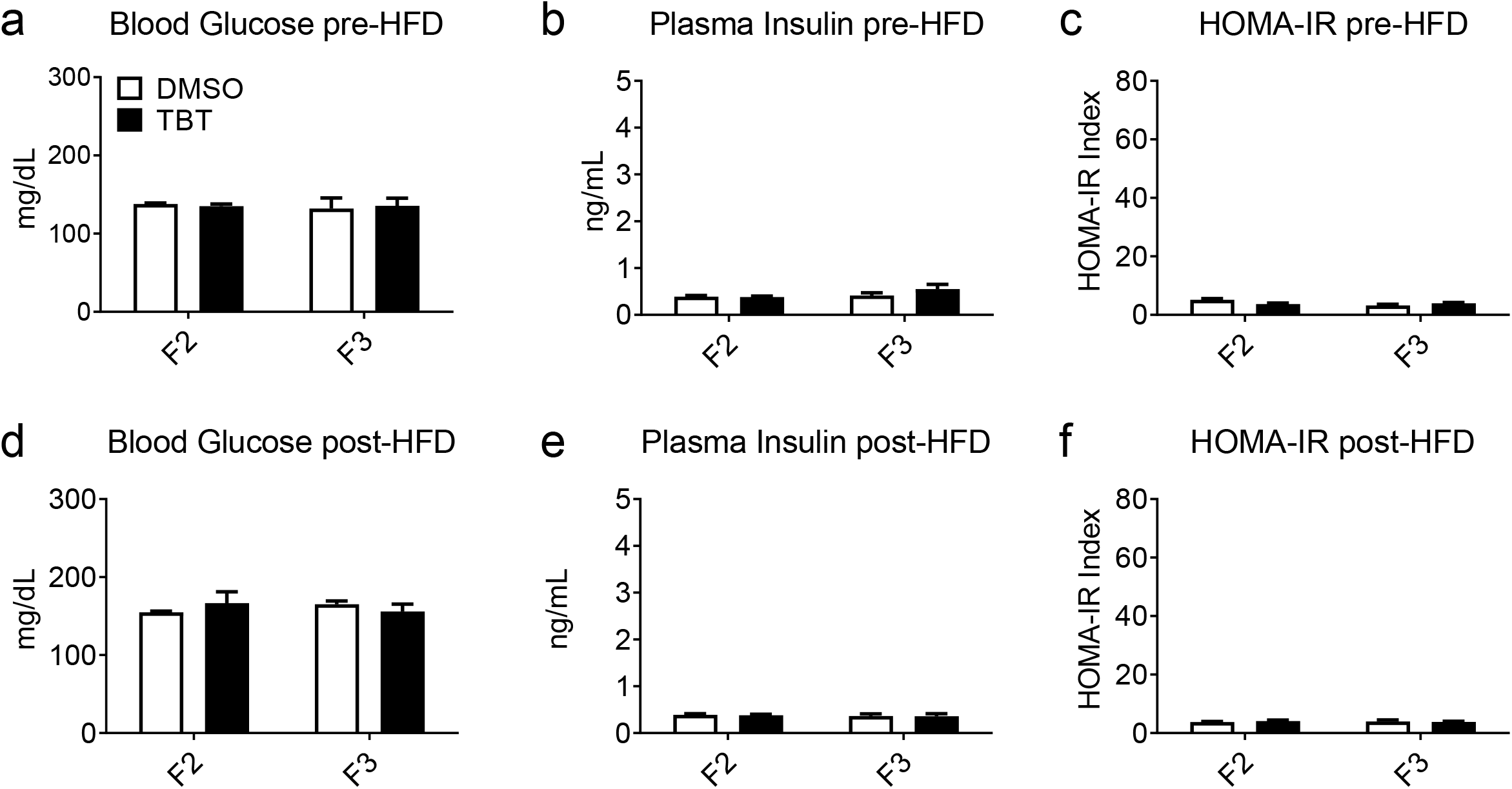
Insulin sensitivity parameters in female descendants. **a**, Plasma glucose and **b**, insulin of female mice before diet challenge (F2 at 5 weeks age and F3 at 17 weeks age) were measured and **c**, HOMA-IR indexes were calculated accordingly. **d**, Plasma glucose and **e**, insulin of female mice after six weeks of diet challenge (F2 at 11 weeks age and F3 at 22 weeks age) were measured and **f**, HOMA-IR indexes were calculated accordingly. Statistical significance was determined using two-way ANOVA. Pair-wise Bonferroni post-tests were used to compare different groups.

**Extended data Fig. 8.**
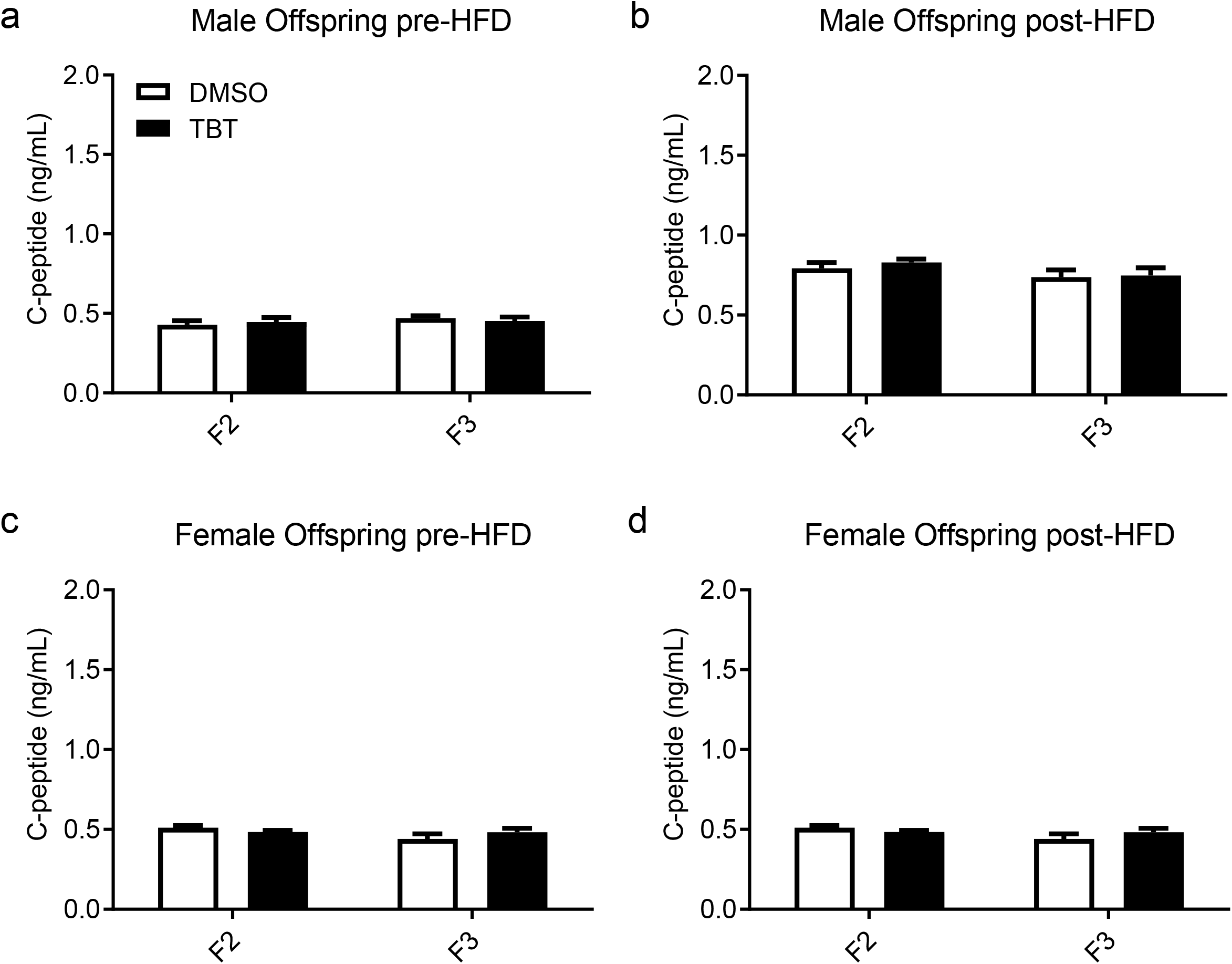
Plasma C-peptide levels in F2 and F3 descendants. Plasma C-peptide concentrations (F2 at 5 weeks age and F3 at 17 weeks age) were measured. **a**, male mice before and **b**, after diet challenge **c**, female mice before and **d**, after diet challenge. Statistical significance was determined using two-way ANOVA. Pair-wise Bonferroni post-tests were used to compare different groups.

**Extended data Fig. 9.**
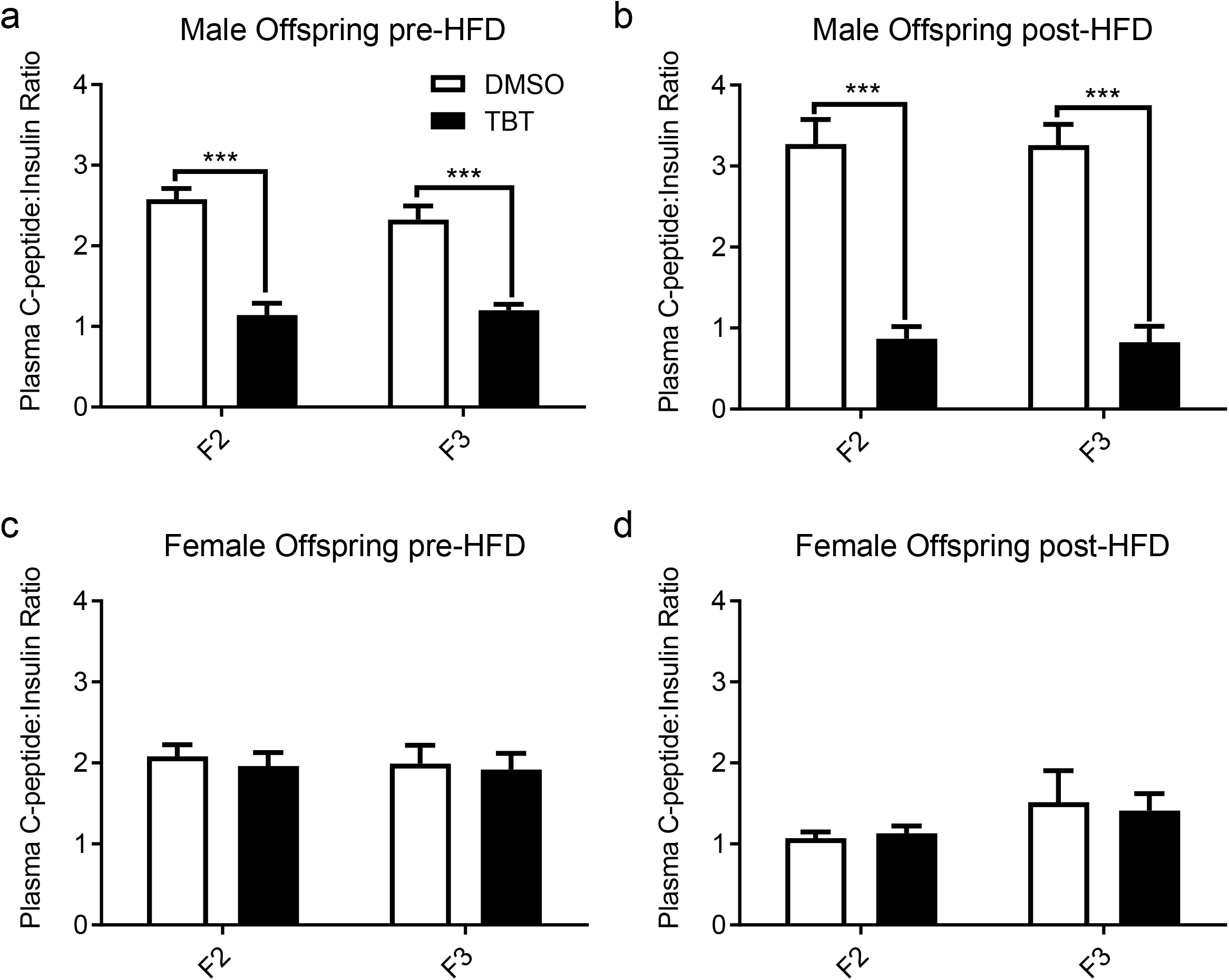
Plasma C-peptide to insulin ratio in F2 and F3 descendants. Plasma C-peptide and insulin of (measured in moles/L) were measured and the ratio of C-peptide:insulin calculated and presented before (F2 at 5 weeks, F3 at 17 weeks) and after (F2 at 11 weeks, F3 at 22 weeks) HFD diet challenge. **a**, male mice before and **b**, after diet challenge. **c**, female mice before and **d**, after diet challenge. Statistical significance was determined using two-way ANOVA. Pair-wise Bonferroni post-tests were used to compare different groups. Data are presented as mean ± s.e.m. *p < 0.05; **p < 0.01; *** p-value < 0.001.

**Extended data Fig. 10.**
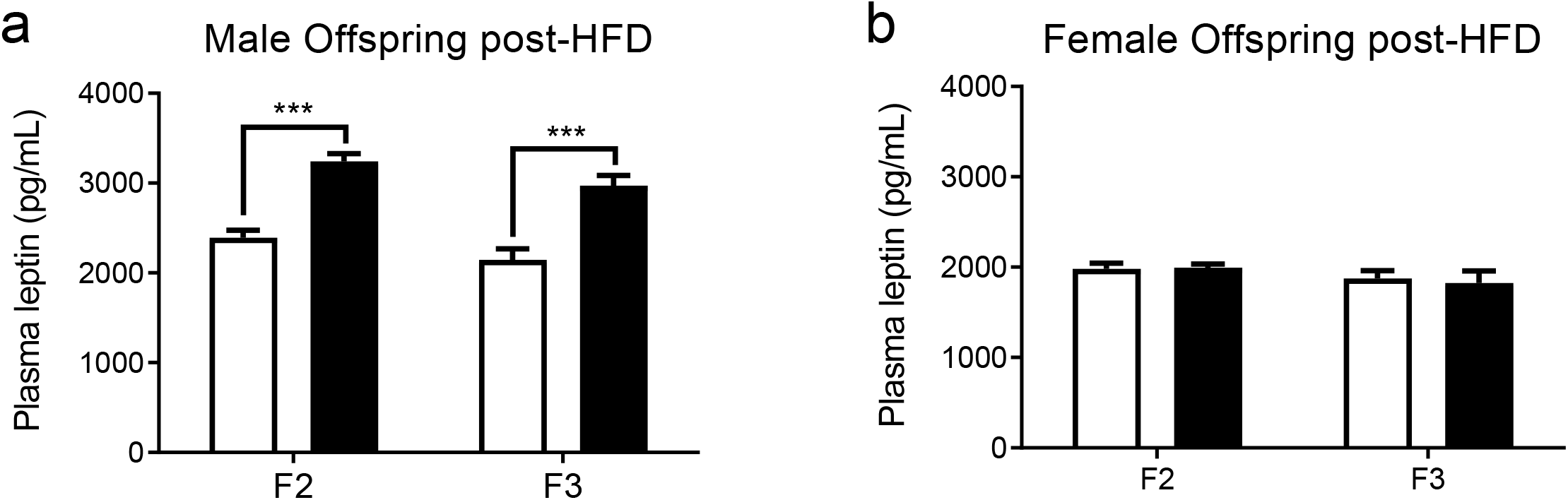
Plasma leptin levels in F2 and F3 descendants. Plasma leptin concentrations of **a**, male mice and **b**, female mice after diet challenge (F2 at 11 weeks age and F3 at 22 weeks age) were measured. Statistical significance was determined using two-way ANOVA. Pair-wise Bonferroni post-tests were used to compare different groups. Data are presented as mean ± s.e.m. *p < 0.05; **p < 0.01; *** p-value < 0.001.

**Extended data Fig. 11.**
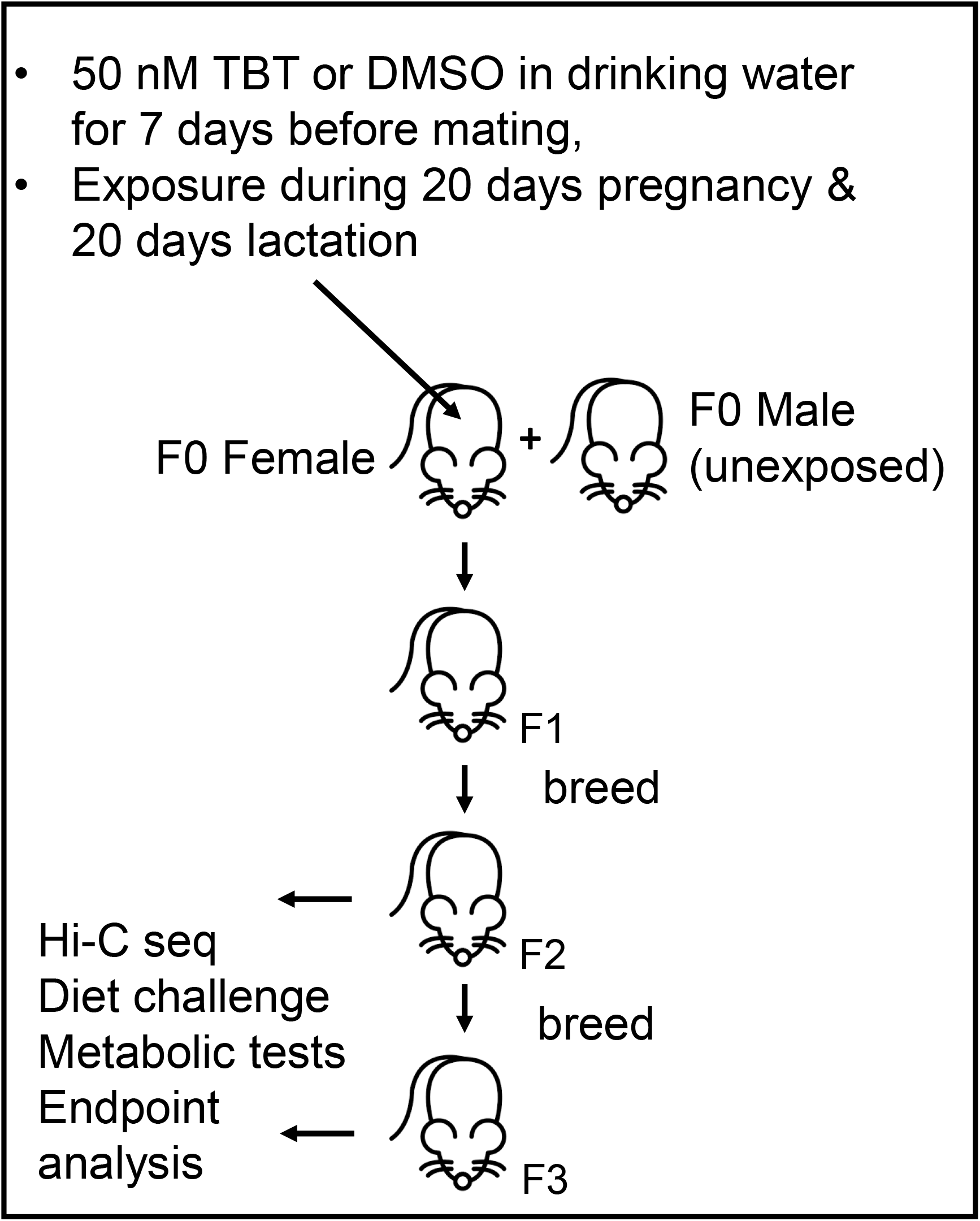
Scheme for breeding, exposure and endpoint analysis. C57BL/6J mice were used in these experiments. Only F0 females were exposed to TBT or vehicle control.

**Extended data Fig. 12.**
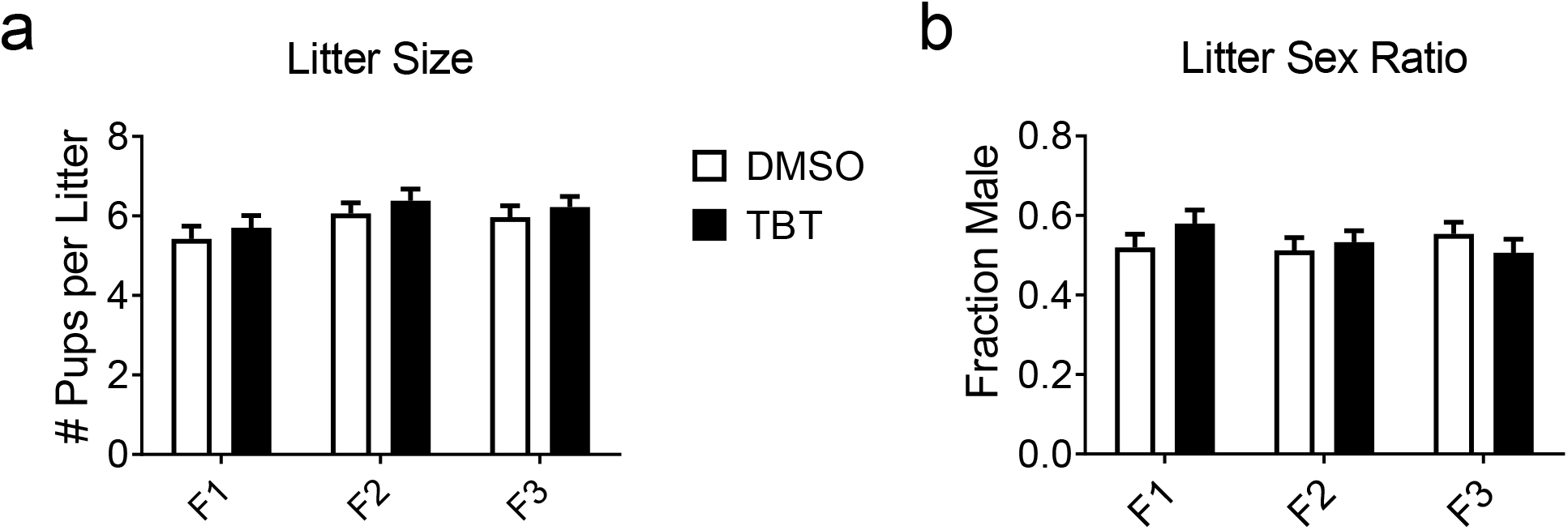
Mice ancestrally exposed to TBT showed no effects on litter size and litter sex ratio of descendants. **a**, Litter sizes of each dam were counted on postnatal day 5, and **b**, sex was distinguished by anogenital distance on postnatal day 14 and confirmed on postnatal day 21 during the weaning processes. 31 to 36 litters of dams from each group were surveyed and data were presented as mean ± s.e.m.

**Extended data Fig. 13.**
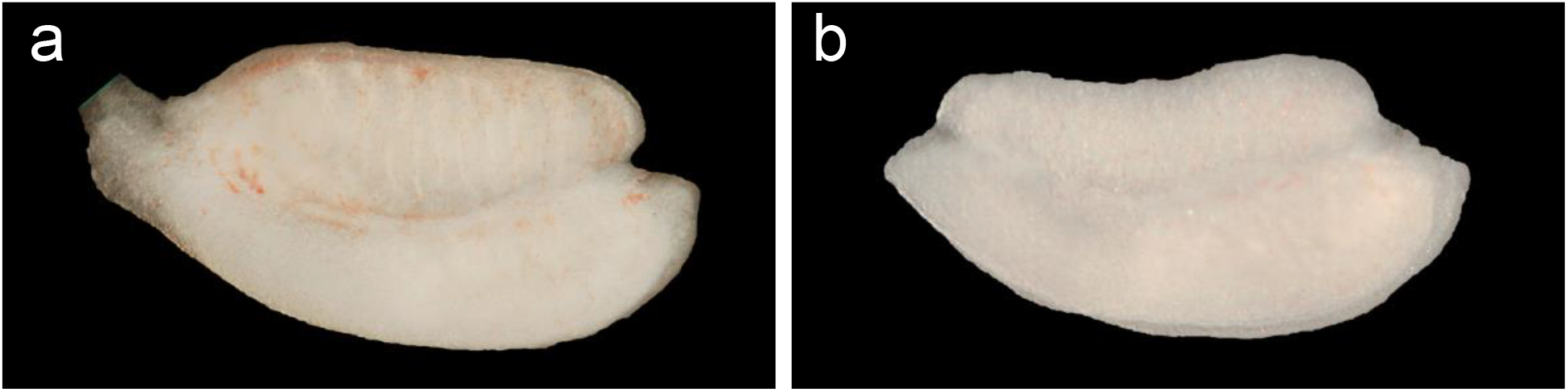
Morphology of dissected gonads. **a**, Male or **b**, female gonads were identified and sexed by their distinguishable characteristic morphology at E13.5. Sex was verified by PCR analysis of the placentas from each embryo prior to ‘omic analysis.

**Extended data Fig. 14.**
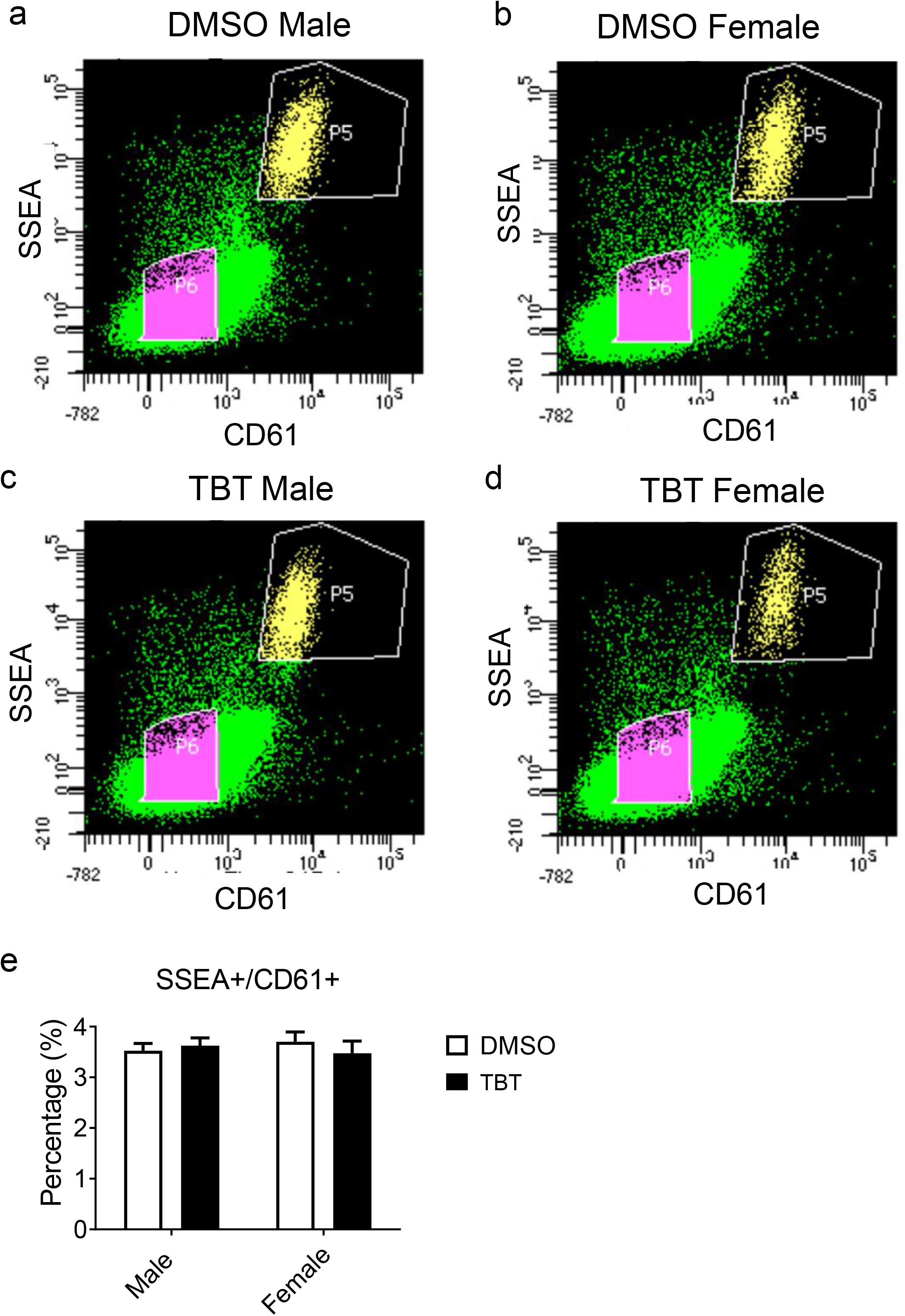
Gating strategy and purity of isolated primordial germ cells (PGCs). Single cell suspension from the enzymatically-digested gonads of **a**, DMSO male descendants, **b**, DMSO female descendants, **c**, TBT male descendants, and **d**, TBT female descendants were analyzed by BD FACSAria II Cell Sorter. SSEA-1+/CD61+ double positive population were gated as P5 indicating PGCs while SSEA-1-/CD61-double negative population were gated as P6 indicating gonadal somatic cells (GSCs). Percentages of PGCs compared to gonad numbers were plot and presented as mean ± s.e.m from 10 litters of each group. Statistical significance was determined using two-way ANOVA. Pair-wise Bonferroni post-tests were used to compare different groups.

**Extended data Table 1.**
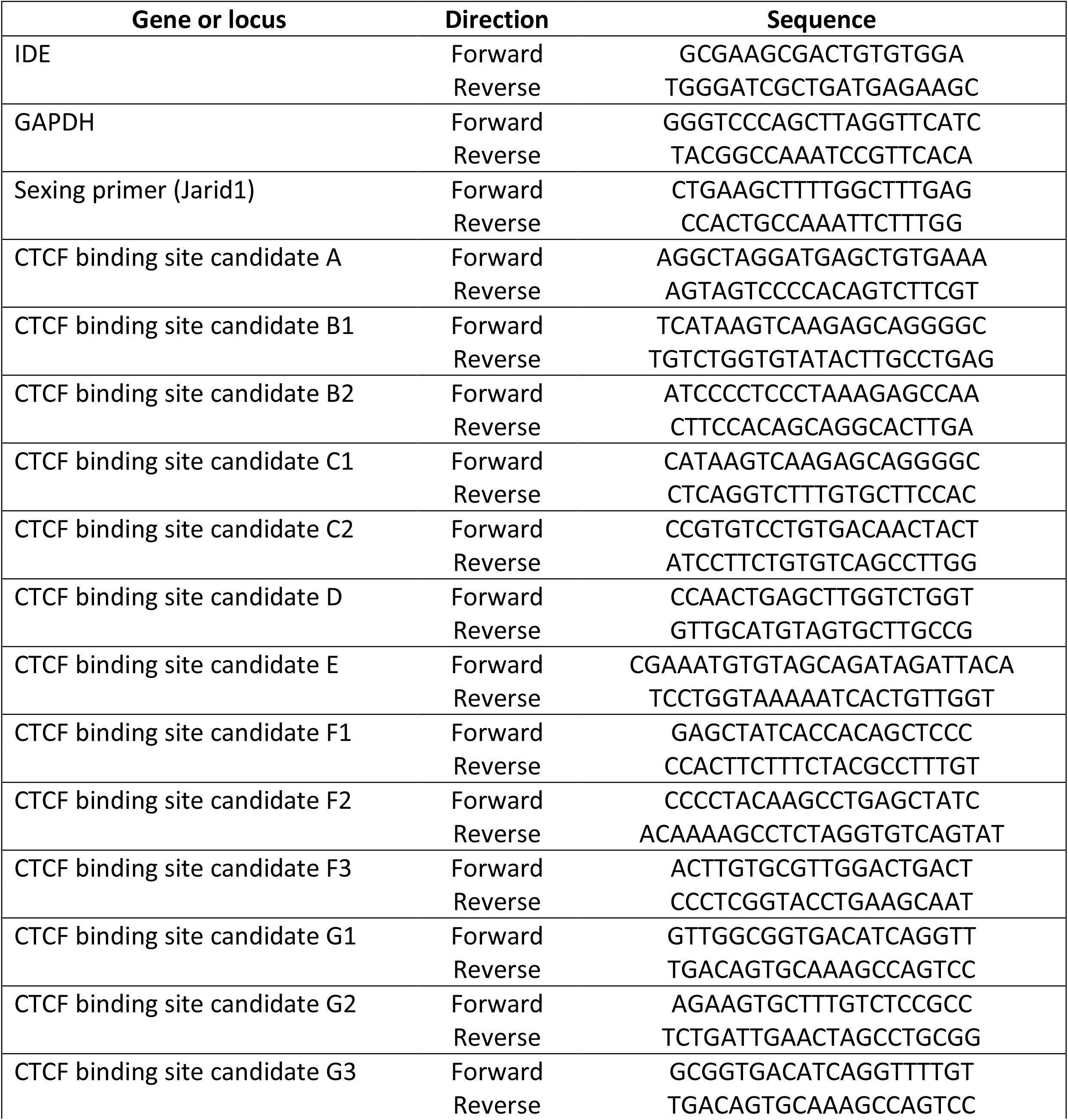
sequences of qPCR primers used in this study.

